# Combinatorial quorum sensing in *Pseudomonas aeruginosa* allows for novel cheating strategies

**DOI:** 10.1101/313502

**Authors:** James Gurney, Sheyda Azimi, Sam P. Brown, Stephen P. Diggle

## Abstract

In the opportunistic pathogen *Pseudomonas aeruginosa*, quorum sensing (QS) is a social trait that is exploitable by non-cooperating cheats. Previously it has been shown that by linking QS to the production of both public and private goods, cheats can be prevented from invading populations of cooperators and this has been termed ‘a metabolic incentive to cooperate’. We hypothesized *P. aeruginosa* could evolve novel cheating strategies to circumvent private goods metabolism by rewiring its combinatorial response to two QS signals (3O-C12-HSL and C4-HSL). We performed a selection experiment that cycled *P. aeruginosa* between public and private goods growth media and evolved an isolate which rewired its control of cooperative protease expression from a synergistic (AND-gate) response to dual signal input, to a 3O-C12-HSL only response. We show that this isolate circumvents metabolic incentives to cooperate and acts as a combinatorial signaling cheat, with a higher fitness in competition with its ancestor. Our results show three important principles; first, combinatorial QS allows for diverse social strategies to emerge; second, that restrictions levied by private goods are not sufficient to explain the maintenance of cooperation in natural populations and third that modifying combinatorial QS responses could result in important physiological outcomes in bacterial populations.

## INTRODUCTION

Social traits and sociality in bacteria have received extensive attention in recent years, from the exploitation of collective behaviors by cheats [1-10], to the implications for infection [6, 11, 12]. *Pseudomonas aeruginosa lasR* quorum sensing (QS) mutants have previously been shown to act as social cheats in environments where QS is required for growth. This is because cheater cells have a higher relative fitness than wildtype strains in mixed populations when QS is required for maximum fitness because they exploit QS-dependent exoproducts produced by wildtype cells [1, 3, 8, 13, 14]. QS *lasR* mutants are also frequently isolated from cystic fibrosis (CF) lungs [14, 15], and they readily arise during long-term selection experiments where the QS controlled production of public goods can be easily exploited [3, 16]. More recently, QS signals themselves have been shown to be exploitable by *lasI* signal cheats in specific environments [7]. A general overview of QS is provided in this recent review [17].

Given the ease by which QS cheats can evolve and spread, solutions must exist to maintain QS in natural populations. Kin selection theory states that social behaviors can be favored if the fitness benefits preferentially promote the survival of individuals sharing those traits [18-20]. Kin selection has been shown to be an important factor in maintaining cooperative behaviors, including QS, in microbes [1, 14, 16, 19, 21, 22]. Dandekar *et al*. demonstrated that by linking QS to both public and private goods (exoprotease production and adenosine metabolism respectively), *lasR* cheats can be prevented from enriching in co-culture with wildtype cells, as ‘cheats’ suffer a loss of direct fitness due to their failure to exploit private goods. This mechanism for restricting the spread of QS cheats was termed ‘a metabolic incentive to cooperate’ [5]. More broadly, this can be viewed as an example of pleiotropy stabilizing cooperation [23].

We tested whether bacterial cells can utilize novel and more complex forms of cheating to exploit loopholes in QS-dependent private goods metabolism, as suggested by our recent work on the functional roles of multi-signal QS systems [24]. *P. aeruginosa* has a complex QS regulatory architecture, featuring behavioral responses to multiple signal inputs. In previous work, we proposed that by using combinatorial (non-additive) responses to multiple signals with different environmental half-lives, individual cells are better able to resolve both social (density) and physical (flow rates/containment) dimensions of their environment. It was predicted and subsequently observed that this synergistic control controls the expression of secreted proteins [24].

Our earlier predictions were made under the assumption of clonal exploitation of a local environment, i.e., adaptation in the absence of cheats. However combinatorial signaling opens the door to novel social strategies, by tuning the extent of production and response to combinations of signals [24]. Tuning combinatorial responses can in principle decouple the regulation of traits that are individually advantageous (e.g., adenosine metabolism) from traits that are collectively beneficial (e.g. secretion of a costly digestive exo-enzyme). For example, if a cheat was capable of responding to just one signal to control a privately beneficial trait, then losing a response to a combination of QS signals could result in the cheat being able to do several things: (1) survive restrictions levied by private goods metabolism as the cell is still capable of activating metabolism by responding to the single signal; (2) exploit the production of public goods by cooperators by producing lower amounts of public goods, because cooperators responding to multiple signals will produce a greater amount than the cheat. Therefore, combinatorial cheats would potentially enrich at the expense of wildtype cells responding synergistically to multiple signals [24].

We performed a selection experiment explicitly designed to test whether cells can evolve to exploit synergistically-regulated public goods cooperation, given a metabolic incentive to cooperate. We cycled a *P. aeruginosa* double QS signal mutant (*ΔlasI/rhlI*) between growth in private and public goods media, with defined concentrations of synthetic signals. Having a double synthase mutant allowed us to control the signal concentration and test the combinatorial response to experimentally defined signal environments. Cycling the media allowed for a selection pressure on the private good metabolism to be maintained (in adenosine) while allowing the evolution of the public good-dependent responses (Bovine Serum Albumin, BSA). We show that (1) an isolate emerged that can circumvent metabolic incentives to cooperate; (2) this isolate acts as a cheat and has a higher fitness in competition with its ancestor and (3) disruption of combinatorial signaling was not directly linked to mutations in the known QS cascade of *P. aeruginosa*. Our findings demonstrate that combinatorial sensing allows for the evolution of novel cheating strategies, highlighting that bacteria can exploit social behaviors in a number of different ways.

## RESULTS

### Combinatorial signaling determines *P. aeruginosa* fitness and QS-dependent gene expression in a QS-dependent environment

We first tested the impact of combinatorial signaling on strain fitness in an environment where QS is required for maximal growth (QSM) [13]. We used a *PAO1ΔlasI/rhlI* mutant strain (PAO-JG2) that makes no AHL signals, but which contains a *lasB::luxCDABE* chromosomal reporter fusion for monitoring QS-dependent gene expression. We measured the changes in *lasB* expression and PAO-JG2 growth in QSM with no added signals; C4-HSL or 3O-C12-HSL added in isolation; or a combination of both signals. We found using OD_600_ and relative light production, that both the growth of PAO-JG2 (Fig. 1A) and *lasB* expression (Fig. 1B) in QSM, shows a synergistic (‘AND-gate’) combinatorial response. i.e. the response to dual signal inputs was greater than the additive combination of responses to single signal inputs alone (χ^2^ = 8.3 *p* = 0.003 and χ^2^ = 6.8 *p* = 0.009 respectively).

**Figure 1.**
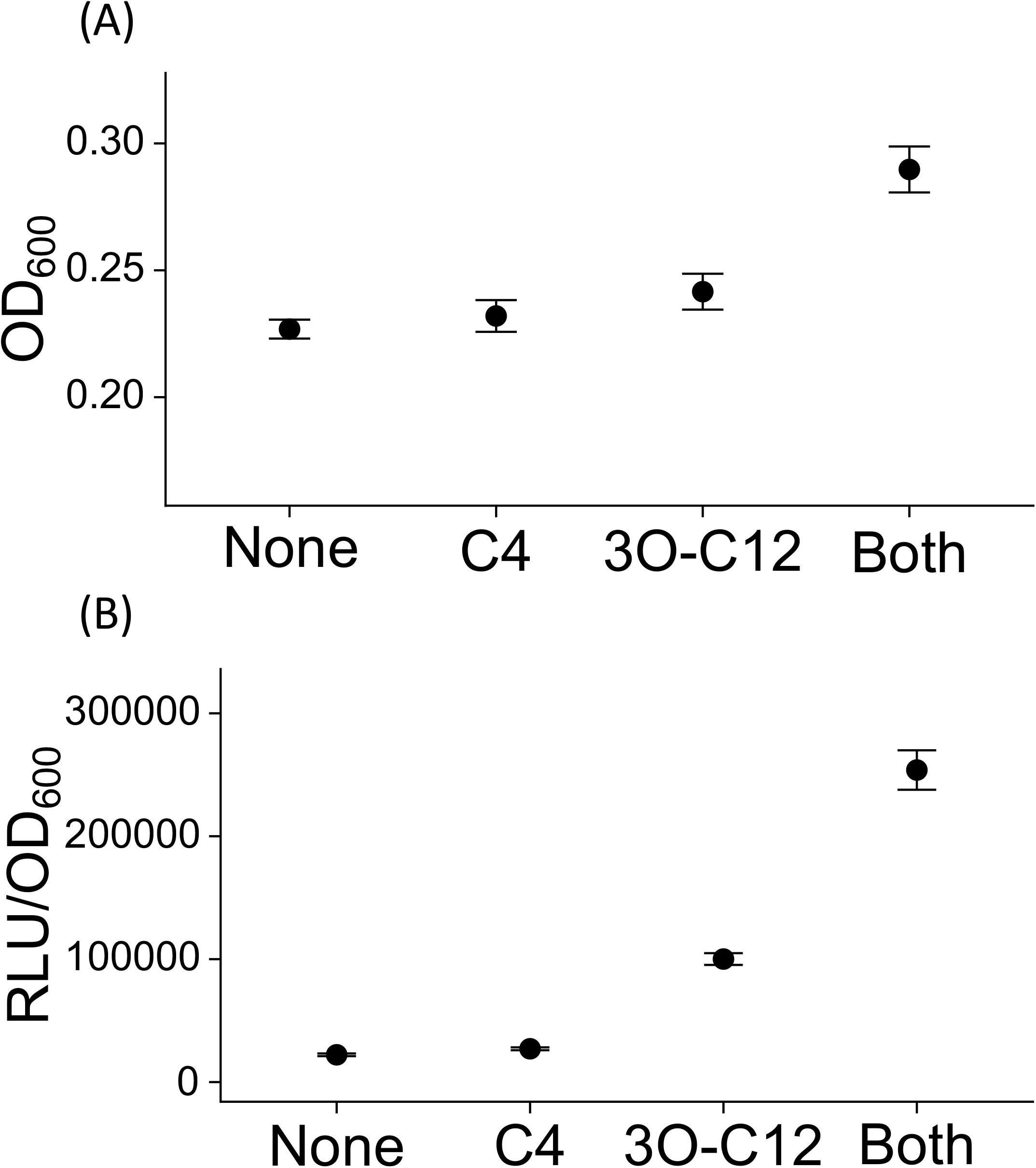
Combinatorial signaling (AND gate) determines *P. aeruginosa* growth and QS-dependent gene expression in QSM. A *P. aeruginosa ΔlasIrhlI* mutant (PAO-JG2) responds in a synergistic manner to both 3O-C12-HSL and C4-HSL signals with respect to (A) growth in QSM; (B) *lasB* expression in QSM. Each signal was added at a concentration of 0.5 μM. Error bars represent ± SEM of 5 replicate experiments.

### Exploitation of combinatorial sensing allows for novel cheating strategies

Combinatorial signaling theory predicts that secreted products (which can be socially exploited) are controlled synergistically by multiple signal molecules, and consistent with this, we find that *lasB* expression is under AND-gate control (Fig. 1A&B) [24]. In contrast, we found previously that intracellular (private) adenosine metabolism by the purine nucleosidase Nuh (*nuh*) is under single signal control (a 3O-C12-HSL gate) [24]. Given these signal processing rules, a simple *lasR* cheat strategy (no response to 3O-C12-HSL) will profit when mixed with wildtype cells in QSM – but will pay a cost if growth is at all dependent on adenosine metabolism. We next asked whether bacteria can overcome this metabolic incentive to cooperate by evolving their signal processing rules to produce novel and more elaborate combinatorial cheat strategies.

To test whether combinatorial cheats can evolve, we performed an evolution experiment (see Materials and Methods for details; Fig. S1). Briefly, our experiment involved cycling the bacteria between a private good-dependent growth media (adenosine) where activation via a 3O-C12-HSL gate is required for growth; and in a public good-dependent growth media (QSM) where activation of the AND-gate positively impacts growth. After 10 rounds of selection we tested the expression of *lasB::lux* in six independently evolved populations in response to 3O-C12-HSL (Fig. S2A). A single population (evolved line C) showed a clear reduction in *lasB* expression levels with the addition of both signals but still responded more than would be expected without any added signal. We tested 5 random individual isolates from this population and each demonstrated the same response to signals as the whole population (Fig. S2B). We selected individual A from line C as our **<u>N</u>**on-**<u>C</u>**ombinatorial **<u>R</u>**esponding mutant (NCRi) that could respond to 3O-C12-HSL but had lost the ability to respond in a combinatorial manner to both 3O-C12-HSL and C4-HSL together. When grown in QSM in monoculture, this mutant did not display any significant increases in growth (Fig. 2A; 3O-C12-HSL vs Both *p* = 0. 540, post-hoc Dunn test) or *lasB* expression (Fig. 2B; 3O-C12-HSL vs Both *p* = 0. 631, post-hoc Dunn test) in QSM in the presence of both signals when compared to the single signal treatment.

**Figure 2.**
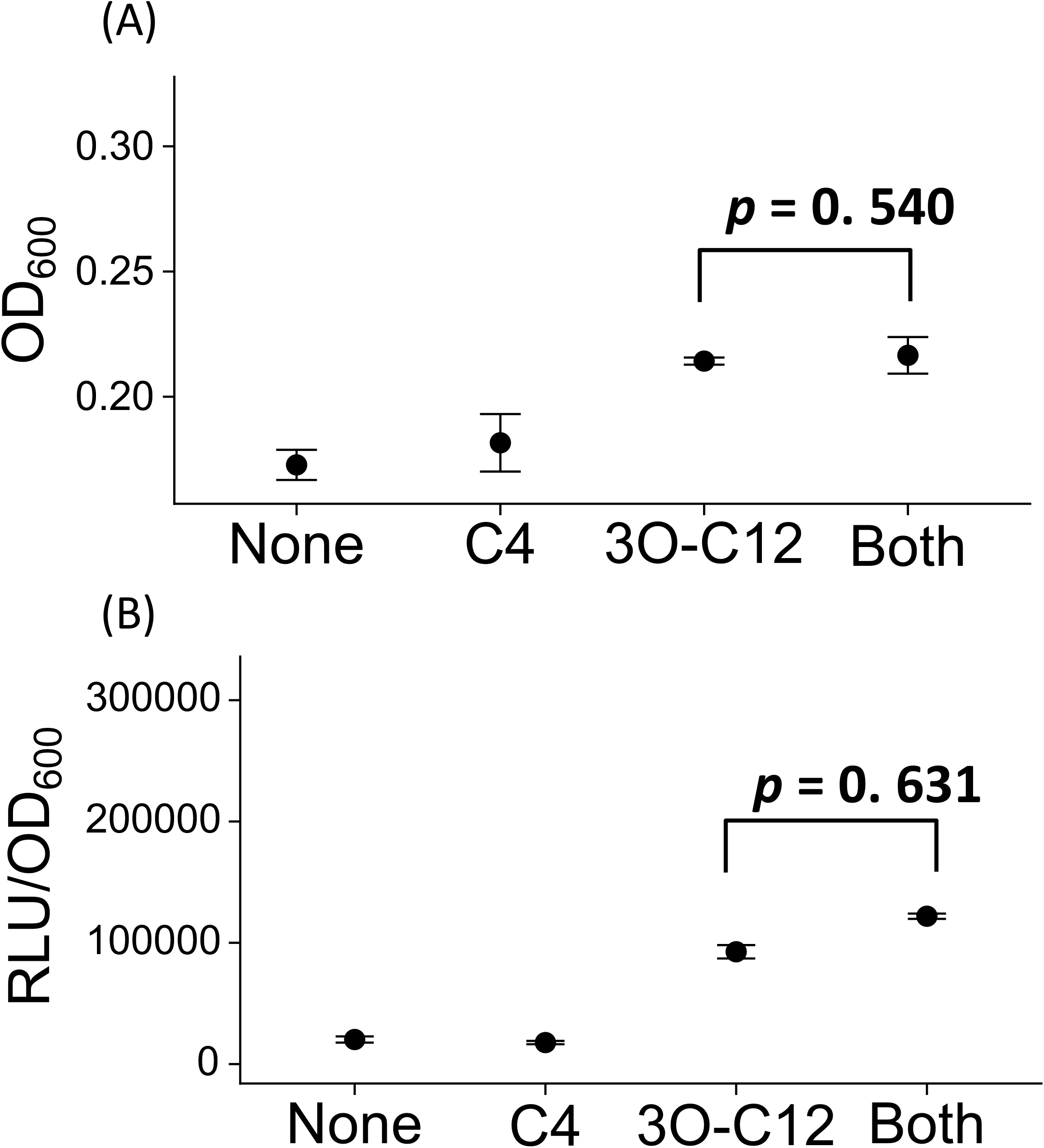
Combinatorial signaling is altered in an evolved isolate. The evolved mutant (NCRi) lost the ability to respond to both signals and neither growth (A) or *lasB* expression (B) was increased in QSM with the addition of both signals. Each signal was added at a concentration of 0.5 μM. Error bars represent ± SEM of 5 replicate experiments.

To test whether the evolved isolate could cheat on a strain that responds synergistically to two signals, we performed a competition experiment. We used an un-labeled PAO1Δ*lasI/rhlI* strain (PAO-JG1) [24] as the competing strain, to be able to distinguish between competing strains in co-culture using light production. We also competed the PAO-JG2 ancestor against PAO-JG1. The only difference between PAO-JG1 and PAO-JG2 is that PAO-JG2 contains a *lasB::luxCDABE* fusion. We found that the fitness of PAO-JG2 compared to PAO-JG1 did not change when either 3O-C12-HSL alone or 3O-C12-HSL and C4-HSL together were added to competition experiments (Fig. 3). An expected result as these strains are essentially the same, with the only difference being the *lasB* reporter in PAO-JG2. We saw no difference in the relative fitness of NCRi compared to PAO-JG1 when 3O-C12-HSL was added in isolation (Fig. 3; *p* = 0.702, Post-hoc Tukey). Under these conditions, both PAO-JG1 and NCRi are capable of growth and express *lasB* to a similar extent (Figs. 1 and 2), and so it is likely that both are contributing public goods. Although it should be noted that NCRi had a reduced growth yield which remains to be explored. In contrast, when both signals were added together, there was a significant increase in the relative fitness of NCRi (Fig. 3; *p* = 0.00015, including the variation of the Grubbs outlier, post-hoc Tukey). Under these conditions, the addition of both signals induces higher growth and *lasB* expression in the ancestor strain compared to NCRi due to a functional response to both signals. In this case, the mutant has reduced QS but still demonstrates increased fitness which is consistent with social cheating. This differs from previous cheating assays, because the NCRi strain is exploiting the synergistic regulation (by multiple QS signals) of public goods production.

**Figure 3.**
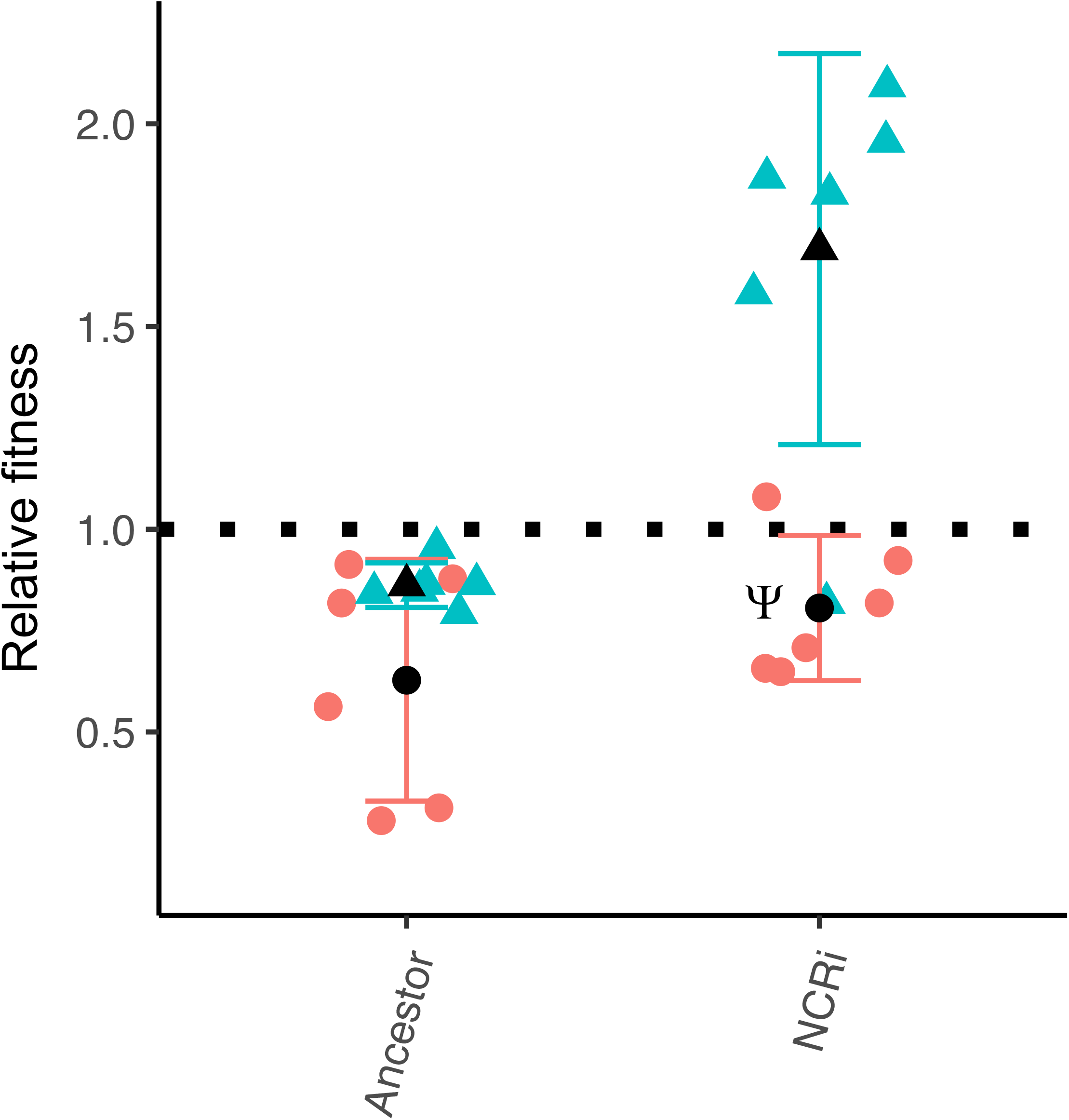
Cheating within combinatorial signaling. The evolved mutant (NCRi) relative fitness is increased when grown with a synergistically responding strain. Both PAO-JG2 (ancestor) and NCRi were competed against a *non-lux* labelled PAO1*ΔlasI/rhlI* strain (PAO-JG1). PAO-JG2 and NCRi each began at a 1:10 frequency with PAO-JG1, with either the addition of 0.5 μM 3O-C12-HSL alone (Red circles) or 0.5 μM 3O-C12-HSL & C4-HSL in combination (Blue triangles). The labelled ancestor strain (PAO-JG1) showed no significant increases in fitness compared to the unlabeled PAO-JG1 in all treatments. The evolved strain NCRi demonstrated a significant increase in relative fitness when both signals were added in combination. Error bars are ± SEM with 6 replicates. The black spot is the mean value. We also identified a Grubbs outlier in the NCRi line with both signals, denoted by (Ψ). Our statistical analysis included this outlier variance as the result were still significant.

### Complementation of *rhlR* did not restore the combinatorial response

Our sequencing of NCRi showed that the major regulators of QS were intact, suggesting possible indirect effects on the QS system. As we found that NCRi lost the ability to respond to C4-HSL but not 3O-C12-HSL, we complemented the strain with a constitutively expressed *rhlR* gene. This had a minimal effect on *lasB::lux* expression and did not enable NCRi to respond to the same synergistical level as the ancestor strain (Fig. 4; post-hoc Tukey Test*p* < 0.0000001).

### The loss of combinatorial response in NCRi is not due to mutations in known QS genes

Whole genome sequencing of NCRi detected no mutations in genes or intergenic regions previously s

**Figure 4.**
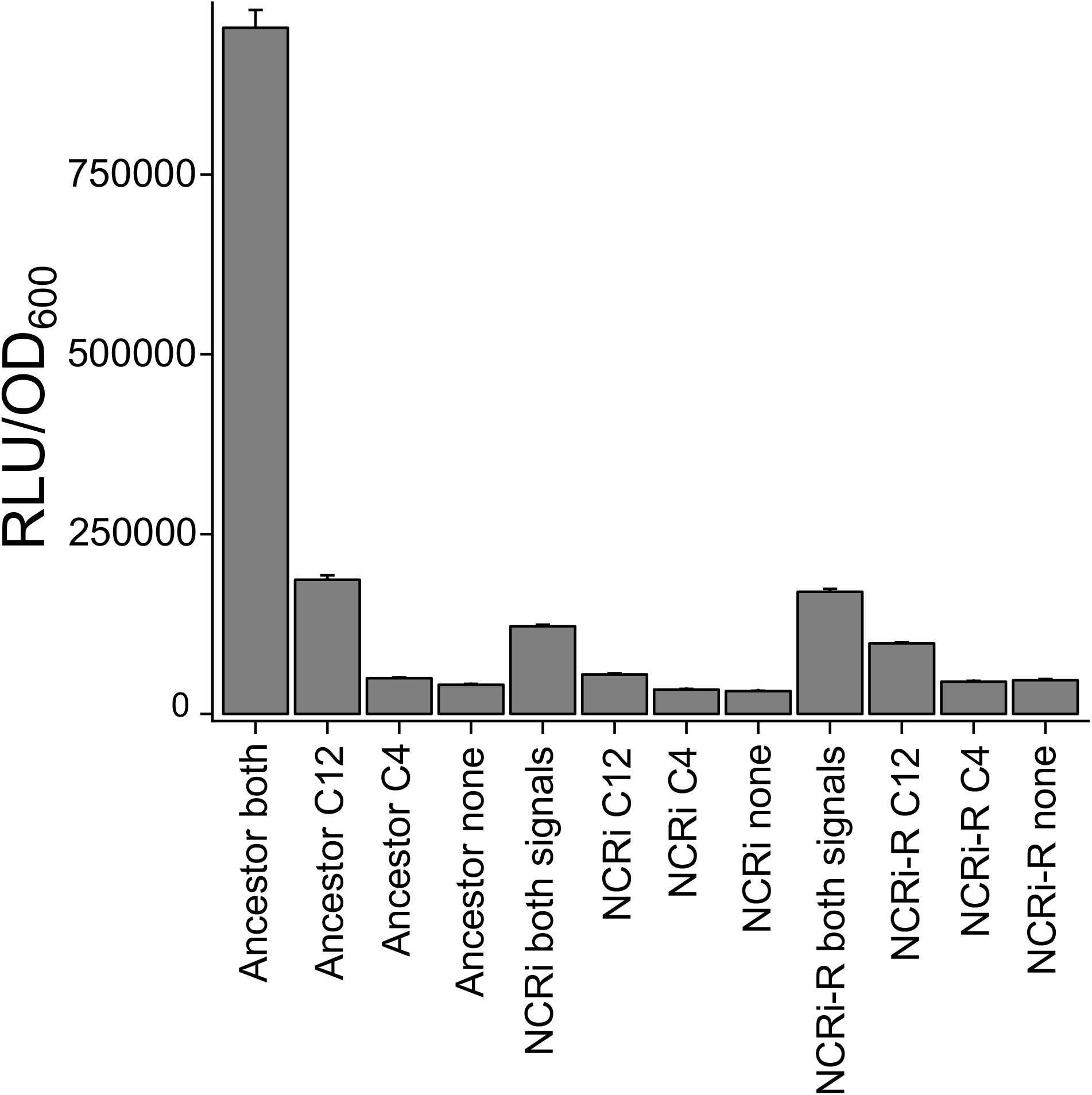
Complementation with RhlR did not restore synergistic response. NCRi was complemented with *rhlR* on a constitutive promoter (NCRi-R) to determine whether indirect effects on *rhlR* were due to lack of synergistic response. Complementation increased the response to all treatments including the control, but a synergistic response was not seen. Signals added at 5 μM each, error bars are ± SEM with 5 replicates.

hown to be involved in the QS-regulatory cascade in *P. aeruginosa*. Importantly, the regulators *lasR, rhlR*, and *pqsR* were all-intact and identical when compared to the ancestor strain and the PAO1 reference library. Four clear mutations were found in NCRi: 2 genes *mexT* and *eftA* had mutations and 2 intergenic regions between *tyrZ* and PA0668 and between PA3504 and PA3505 (Table S1).

## DISCUSSION

A rapidly growing body of research has demonstrated that bacterial QS is a social behavior that is exploitable by cheats [1, 2, 6, 8, 9, 16, 17, 24-26], and a number of explanations for maintaining QS in natural populations have been provided. Kin selection has previously been shown to maintain QS in populations because it increases the reproductive success of producer cells (direct fitness) and other individuals that carry the same QS genes (indirect fitness) [1, 16, 18-20, 27]. The effect of kin selection on QS relies on the fact that QS controls public goods production (such as exoproteases) that can be shared locally between cells [1, 3, 13]. We are now becoming increasingly aware that the QS system does not just regulate the production of public goods, but it also controls private goods metabolism within cells. By linking QS to both public and private goods, it has been shown that this can prevent the invasion of public goods cheats [5, 7, 13, 24]. This has been termed a ‘metabolic incentive to cooperate’ [5].

Here we describe a selection experiment (Fig. S1), specifically designed to test whether metabolic incentives for cooperation can be circumvented by cheats. We based our experimental approach on a number of previous findings. Firstly, that QS-dependent exoprotease production is required for maximal growth in a medium containing a carbon source that is degraded by proteases which act as public goods [1, 3, 13]. Secondly, that expression of the *lasB* gene required for protease (elastase) production is synergistically increased by the addition of C4-HSL and 3O-C12-HSL in combination [24, 28-31]. Thirdly, that the enrichment of *lasR* cheats can be controlled by private goods metabolism such as growth in adenosine [5, 32].

Our findings show that there are fitness benefits in using a combination of C4-HSL and 3O-C12-HSL to regulate public goods that break down a protein carbon source in QSM (Fig 1A). Many factors could impact on the combinatorial response. Shifts in mass transfer, the total level of rich carbon source, the spatial structure of the population, and levels of potential microbial crosstalk, will almost certainly change the benefits of QS fitness of cells responding in a combinatorial manner [13, 24]. Fitness benefits provided by combinatorial signaling, therefore, opens the door for conditional cheats that can access QS-dependent public goods with lower production of metabolically costly proteases.

We identified an evolved isolate (NCRi) that circumvents the QS-dependent metabolic incentives to cooperate. This isolate acts as a social cheat and has a higher fitness in competition with its ancestor, and disruption of combinatorial signaling was not directly linked to mutations in the known QS cascade of *P. aeruginosa*. The response of NCRi to signal treatments showed that it is capable of using 3O-C12-HSL in isolation to make public goods to sequester carbon, but it lost the synergistic (AND-gate) response when exposed to both 3O-C12-HSL and C4-HSL.

We found that this loss of the ancestral synergistic response, resulted in the mutant having a fitness advantage in the presence of both signals (Fig. 3). Although both strains are capable of producing public goods, indicated by the expression of the *lasB::lux* reporter in the presence of 3O-C12-HSL, the mutant lacks the synergistic response, and will therefore produce less public goods. This suggests that in an environment in which QS cooperation is required for maximal fitness, NCRi receives higher fitness benefits when in competition with the ancestor, because individuals that have maintained the synergistic response to the signals will be producing a greater amount of public goods than strains that have rewired their combinatorial responses to respond only to 3O-C12-HSL [1, 3, 13].

A simple mechanistic explanation for the lack of response to C4-HSL would be a defect in the *rhl* QS system; however, when we whole genome sequenced NCRi, we found no changes in the *rhl* genes or a range of known QS genes (including *pqs* genes) that could easily explain our observed phenotype. Whole genome sequencing of NCRi revealed genomic changes (Table S1). First, we observed a mutation in *mexT* which has previously been linked to similar QS phenotypes [33, 34]. It was previously shown that the rewiring of QS circuits such that RhlIR functions in a LasR-null background is caused by inactivation of MexT. It was also demonstrated that it is unlikely that MexT directly affects the expression of *rhlR* or *rhlI*, because neither of these genes contains the conserved MexT binding motif in the promoter region [33]. The *mexT* mutation in NCRi may be conferring a similar QS phenotype, and may be important for the synergistic response to both signals. Further work is required to fully elucidate the regulatory role of MexT in combinatorial QS.

The potential roles of the other 3 identified mutations (*etfA* and the intergenic regions between *tyrZ*-PA0668 and PA3504-PA3505) will require future work to elucidate what functional role these mutations have (if any) on the observed phenotypes. However, a combination of the observed mutations could impact on combinatorial signaling. Overall, our findings suggest novel ways to circumvent and cheat the known QS system in *P. aeruginosa*, which allows strains to evolve to bypass the restrictions levied by QS-private goods metabolism.

The linking of private and public goods production has been suggested as a way of policing QS cheats [5]. More broadly, this is an example of pleiotropy stabilizing cooperative phenotypes [23]. Recent work by dos Santos (which cited the preprint version of this manuscript as supporting evidence), has shown that only under strict criteria where genetic architecture is fixed, does pleiotropy maintain cooperation. Our work demonstrates that genetic architecture can change, and these changes can reduce the linkage of private goods to police public goods as was predicted by dos Santos [35]. Further, the findings presented here and work by others, suggests that many selective pressures work together to control cheats in natural environments. Factors such as migration, spatial structure, regulatory responses and private goods metabolism could all function together to reduce the relative fitness of cheats [24]. Our work shows that the incentives to cooperate placed by metabolic sources are not adequate to explain the persistence of cooperation in *P. aeruginosa*. Combinatorial responses allow for a greater range of social strategies to emerge than previously considered. Further our results should be taken as preliminary evidence that there remain novel regulators of QS in *P. aeruginosa*, that are capable of regulating protease production and presumably other QS-controlled genes.

## MATERIALS AND METHODS

### Bacterial strains and growth conditions

We used three strains in this study, and we specifically generated PAO-JG2 for the work. This strain is generated from PAO-JG1, a double QS signal synthase insertion mutant (*ΔlasI/rhlI)* [24]. We integrated a *lasB::luxCDABE* fusion constructed using the mini-CTX::*lux* system [36] into the chromosome of PAO-JG1 to create PAO-JG2. We also used PAO-JG1 as an unlabeled *lux* strain for competition assays. The other strain we used in this study is NCRi, a mutant of PAO-JG2, generated in an evolution selection experiment (see below). We routinely grew liquid cultures of strains at 37 °C with shaking (200 rpm) in 5 ml of modified Quorum Sensing Medium (QSM) [13]. We monitored the growth of bacterial cultures by measuring the absorbance of light at a wavelength of 600 nm (optical density, OD_600_) using a spectrophotometer (Tecan 200i). We supplemented M9 agar plates with 0.4% adenosine as the sole carbon source [5].

### Selection experiment

To test whether bacteria can avoid restrictions imposed by the metabolism of private goods by exploiting combinatorial signaling, we designed an experimental evolution approach (Fig. S1). Our experimental protocol (by design) prevented the *de novo* generation and spread of *lasR* mutants, but allowed for mutants that could exploit combinatorial signaling to evolve and avoid the levies imposed by QS-dependent private goods metabolism. Our selection experiment was performed in 6 replicate lines (each containing 4 microcosms) using PAO-JG2 as the ancestor strain, and we cycled evolving populations between QSM [1, 13] and an M9 agar media supplemented with adenosine as the sole carbon source (Fig. S1). We grew PAO-JG2 for 48 h in QSM media at 37°C/200 rpm, and we supplemented the media with 1 μM of each C4-HSL and 3O-C12-HSL. We then plated the 4 microcosms of each replicate line onto adenosine M9 agar plates for 48 h. This plating restricts the passage of *lasR* mutants as they are unable to grow with adenosine as the sole carbon source. As we wished to study novel forms of cheating, we removed *lasR* mutants. We selected single colonies from the plates for each of the 4 new microcosms which we used to seed the next round of selection in QSM media. We ran the selection experiment 10 rounds (a single round being the cycle between growth in liquid QSM and on adenosine plates), which equates to ≈ 70 generations. After each round, we stored whole selected populations in QSM containing 25% glycerol −80°C. After 10 rounds of selection, we tested populations and individual isolates for altered responses to QS signals (*lasB::lux* expression as shown in Fig. S2). We standardized *lasB::lux* measurements by dividing relative light units (RLU) by optical density (OD_600_), resulting in per cell average *lasB* expression. We used peak level responses for analysis, and all peaks fell between 7-10 hours after initiation of growth. We randomly selected a single individual isolate from the final population of replicate line C and found that it showed altered *lasB::lux* expression compared to the ancestor when both signals were added in isolation or combination.

### Competition experiments

We competed the ancestor (PAO-JG2), or our evolved isolate (NCRi), with an unlabeled version of the ancestor strain (PAO-JG1). We started cultures at equal densities (OD_600_ from each was measured then equally partitioned to a final OD_600_ of 0.05), in QSM media ± Signals (3 treatments, no signal, 3O-C12-HSL, and both C4-HSL and 3O-C12-HSL each at 0.5 μM) for 18 h. We then diluted cultures to 10^-6^ CFUs and we plated them out onto LB agar plates. We counted the total number of colonies expressing light (using a light amplifying camera) and compared this to the total number of colonies. We calculated the relative fitness of strains using the formula w = p_1_(1 - p_0_)/p_0_(1 - p_1_) where po and pi are the proportion of the strain we were testing fitness for in the population before and after incubation respectively [37]. In Fig. 3, a single replicate in the NCRi group with both added signals was in line with the addition of only a single signal. Using a Grubbs test, this replicate was identified as an outlier [38], however, even including the increased variation of the outlier, the power of this experiment is still 93% giving us very high confidence that this result is significant and neither a type I or type II error [39].

### Genomics and sequencing

We prepared genomic DNA from 14 h cultures of PAO-JG2 and the evolved NCRi isolate. We extracted DNA using the Sigma GenElute Bacterial Genomic DNA kit following the manufacturer’s guidelines. We performed multiplexed, 150 bp Paired-end sequencing on the Illumina HiSeq3000 platform to an average depth of 150x coverage. We performed de novo assemblies using Spades [40] annotated the assembled genomes using Prokka [41]. To investigate any large scale insertions or deletions between ancestral and evolved strain we performed comparative genomics by pairwise Blast analysis which we visualized using BRIG [42] and by generating a ProgressiveMauve genome alignment [43]. To determine the presence of SNPs in NCRi relative to the ancestral PAO-JG2 strain, we mapped the raw fastq data for both the ancestral PAO-JG2 and NCRi against a reference PAO1 genome using Breseq [44]. The raw sequence data for both strains has been deposited in the SRA under project accession number PRJNA437484.

### Complementation with RhlR

We prepared competent *P. aeruginosa* cells by taking a 1% (v/v) inoculum from an overnight *P. aeruginosa* culture, adding to 50 ml of sterile LB medium and growing at 37°C/200 rpm to reach an OD_600_ of 0.4-0.5. We harvested cells by centrifugation at 5,000 rpm for 10 min at 4°C, and we then washed the cells three times in sterile ice-cold 300 mM (w/v) sucrose solution. We re-suspended cells in 200 μl of the ice-cold sucrose solution and then incubated for 30 minutes on ice. We performed electroporation in 0.2 cm electroporation cuvettes (Flowgen) containing 40 μl of competent cells and 2 μl of purified pUCP18::*rhlR*. We delivered an electroporation pulse of 1.6 kV using a BioRad Gene Pulsar connected to a BioRad pulse controller (BioRad Laboratories, Watford, UK). We then added a 1ml aliquot of LB broth to the cells and incubated for 1 h at 37°C/200 rpm before plating 100 μl onto selective LB agar plates containing carbenicillin (300 μg/ml). We incubated plates overnight at 37°C and selected isolates containing plasmids the next day.

### Statistical analysis

We performed all statistical analyses using R (v3.4.2). We examined population selection lines by either an ANOVA, then applying posthoc Tukey HSD tests. When the data were not normally distributed, we used the Kruskal-Wallis tests within the car package for R. For MIC comparison a Welch two sample t test was performed. The outlier in the competition experiment (Fig. 3) is a Grubbs outlier, we used the Outlier package in R and power was determined with the built in function in R.

## Supplemental information

**Figure S1.**
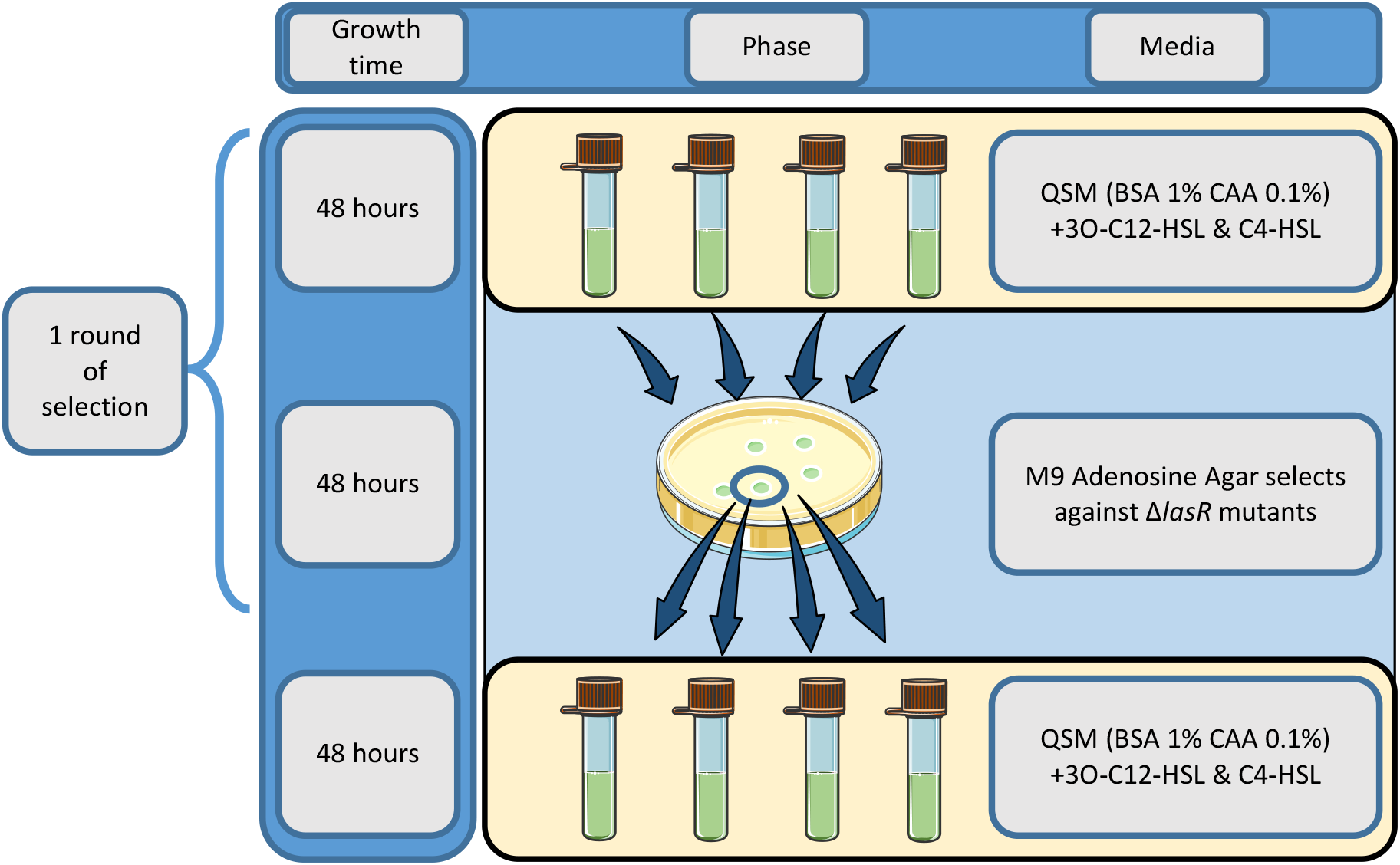
Schematic of selection experiment. Our selection experiment was performed using two separate phases of growth. (1) A liquid phase in which BSA would be broken down by the production of public good enzyme, extracellularly, and thus open to exploitation of individuals producing lower levels of protease. (2) A phase on agar plates where acquisition of carbon was achieved by the intracellular process of adenosine utilization, which requires a functional las QS via 3O-C12-HSL. The selection experiment also used a global selection approach to maintain cooperation. In each replicate line, 4 growth microcosms were pooled at the end of the liquid phase and streaked onto a single adenosine plate, thus promoting the selection of the fittest strains.

**Figure S2.**
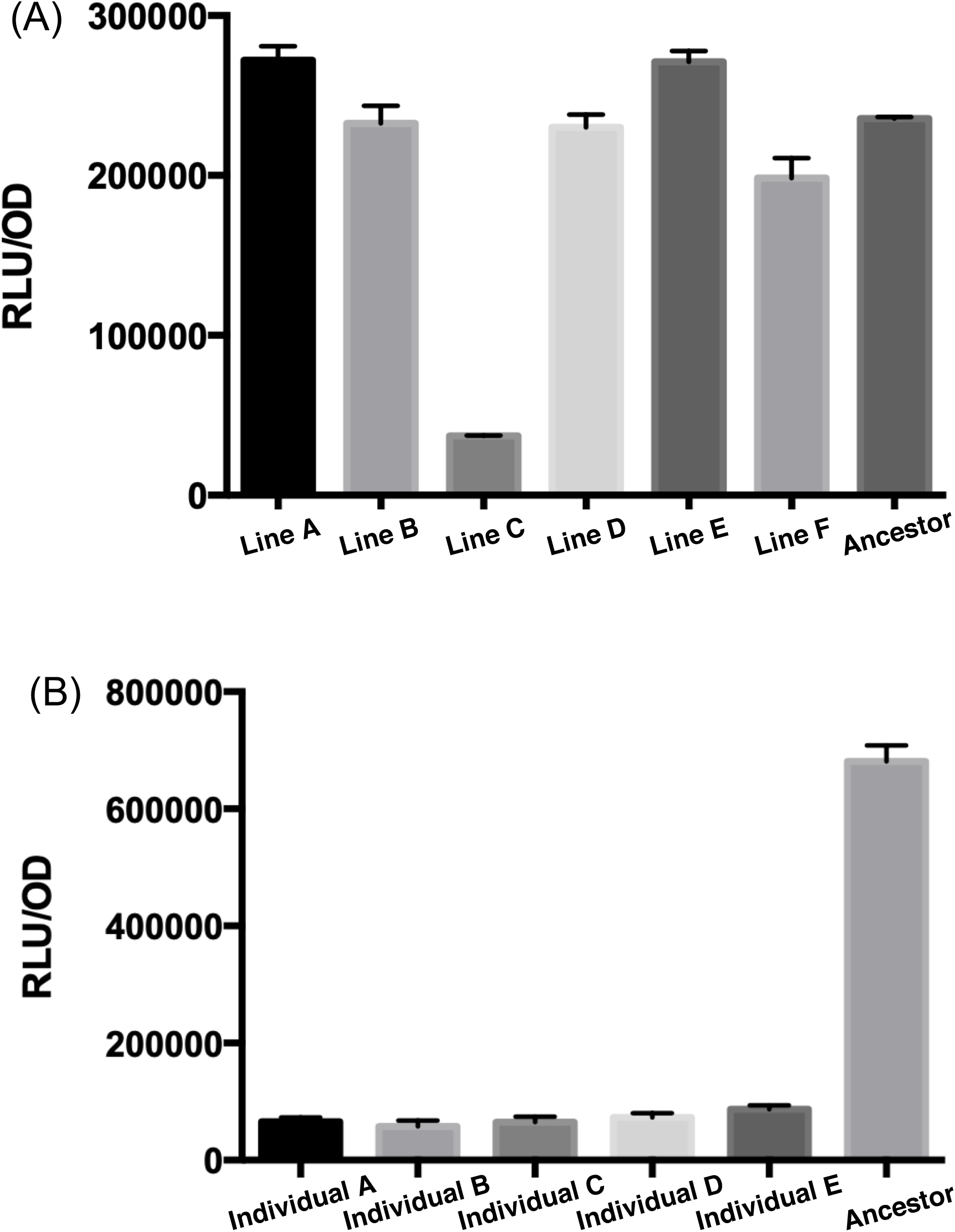
Experimental evolution lines and QS phenotype response. We conducted a large experimental evolution experiment with multiple lines. After roughly 70 generations we examined the QS phenotype of *lasB* expression under the combinatorial response using the *lasB::lux* reporter. (A) Of the 6 replicate lines initiated, a single line (line C) showed a clear reduction in expression level when 5 μM 3O-C12-HSL was administered. (B) We then examined individuals from line C, picking a random set and testing the combinatorial response with 5 μM of each signal and showed that all the picked isolates demonstrated a comparable phenotype. Individual A was picked, stored and named NCRi.

**Table S1.**
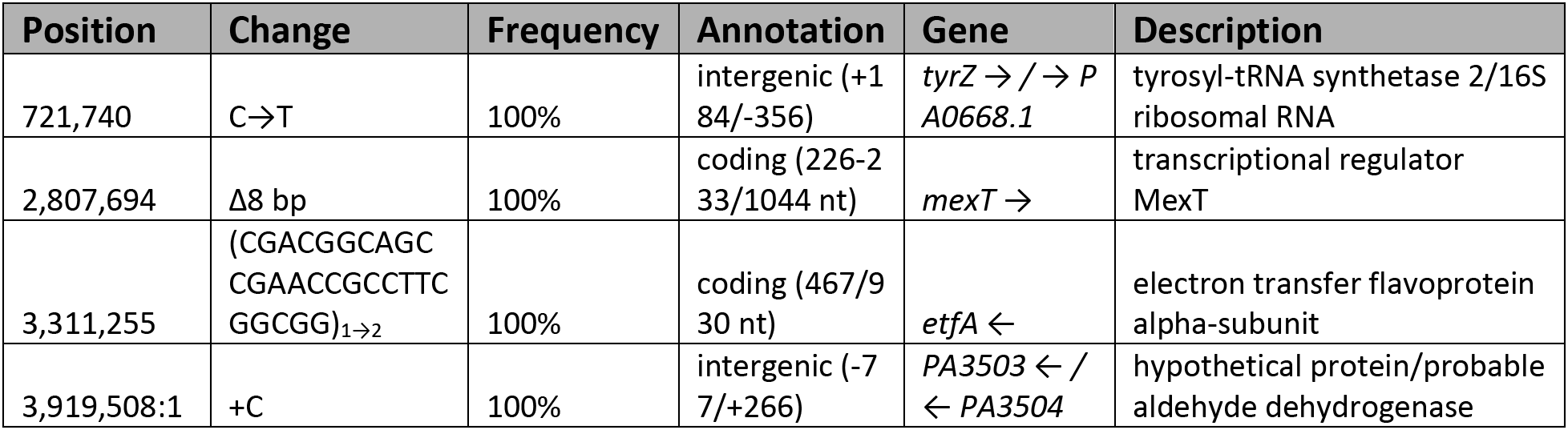
List of Breseq identified mutations in NCRi.

## Acknowledgments

We thank Freya Harrison for help with extracting Genomic DNA and proof checking the statistical analysis and Alan McNally for initial sequencing analysis.

## Author contributions

J.G., S.P.B, and S.P.D. conceived the study; J.G. and S.A. performed the experimental work; J.G. analyzed the data; All authors contributed to writing the manuscript.

